## Supplemental information for "Activation of an injury-associated transient progenitor state in the epicardium is required for zebrafish heart regeneration"

### Supplementary Figures

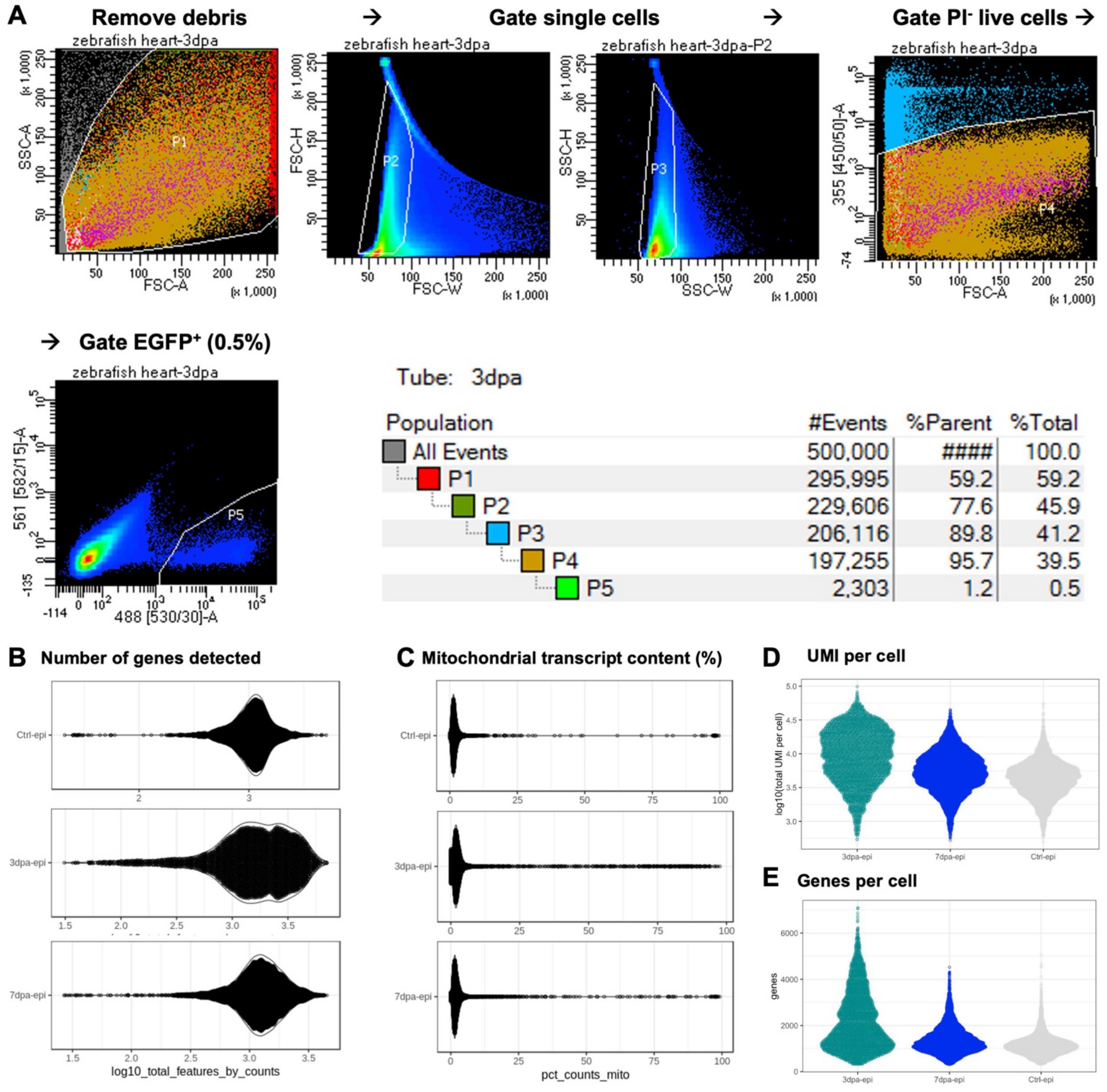


**Figure S1. FACS isolation of single *tcf21*+ cells and QC of scRNAseq.**

(**A**) Flow Cytometry plots showing the cell sorting process. About 0.5% of live ($PI^-$) $GFP^+$ cells were isolated from the ventricular dissociation. (**B**) The numbers of genes detected in each scRNA-seq sample. Each dot represents a single cell; the x-axis depicts the number of genes that were detected within a given cell. Most cells have evidence of at least 1,000 genes. (**C**) Mitochondrial transcript content (% transcripts originating from mitochondrially encoded genes). (**D, E**) Distribution of the UMI content (D) and genes (E) of each sample after removal of low-quality droplets and rarely captured genes. The samples of day 3 post-injury (3dpa) yielded greater numbers of detected genes and, correspondingly, greater numbers of individual transcripts.

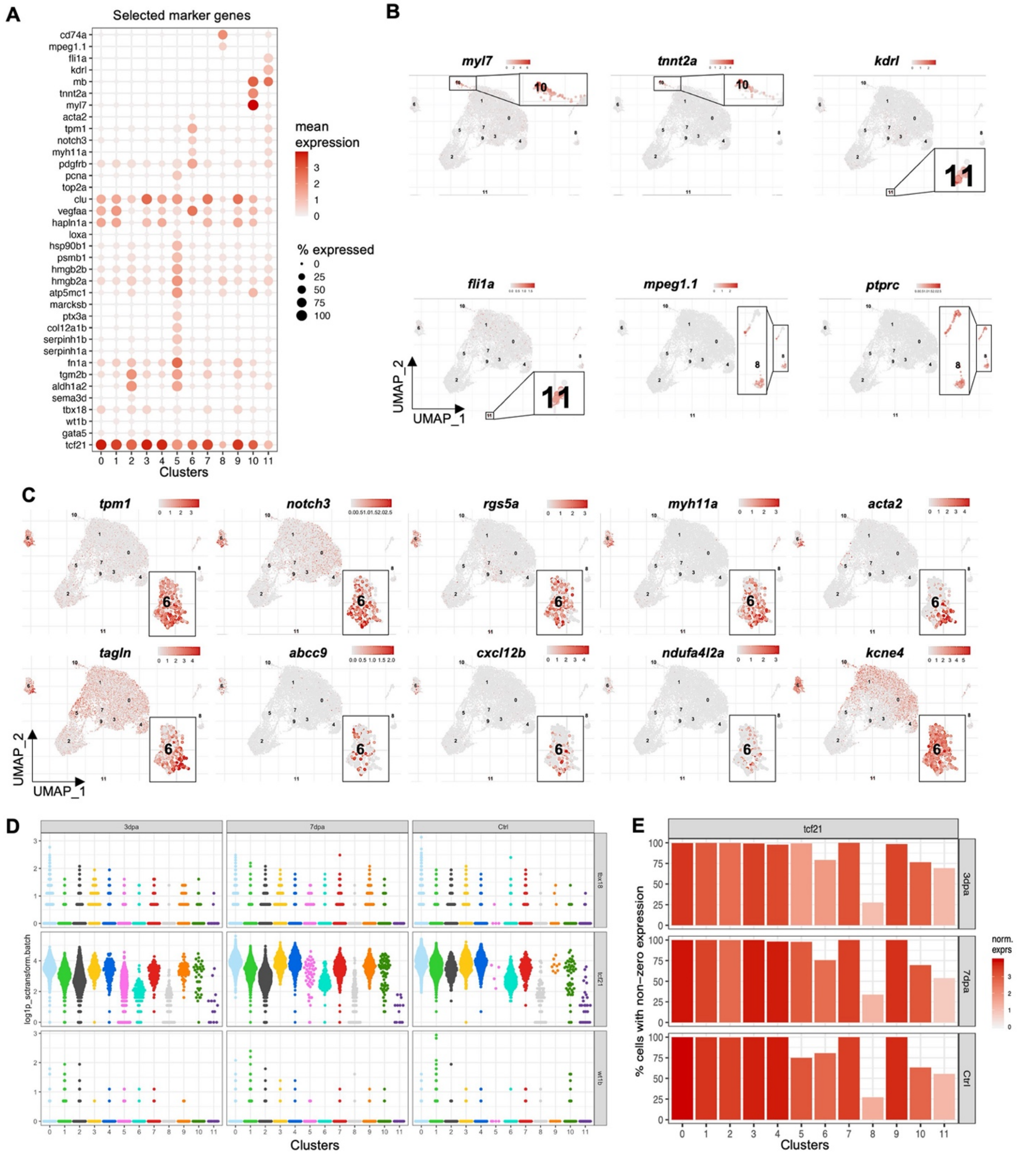


**Figure S2. Selected cluster markers.**

(**A**) Expression of selected cluster marker genes across different clusters. This dot plot depicts the abundance and expression magnitude of individual genes across cells of given clusters. The dot size represents the fraction of cells with at least one UMI of the specified gene. (**B**) Expression patterns of the selected non-epicardial genes within the

UMAP projection (3 samples combined). Magnified views are shown in frames. (**C**) Expression patterns of top mural cell markers on UMAPs (3 samples combined). Magnified views of cluster 6 are shown in frames. (**D**) A jitter plot showing relative expression levels (log1p_sctransform) of *tbx18*, *tcf21*, and *wt1b* in each cell across clusters (0-11) and conditions (Ctrl, 3 dpa, and 7 dpa). (**E**) Visualization of the magnitude and abundance of *tcf21* expression. Each bar's height represents the percentage of cells for which at least one fragment representing *tcf21* was detected. The color corresponds to the mean normalized expression value with dark red, indicating that numerous transcripts of *tcf21* were detected in the cells of a respective cluster (x-axis).

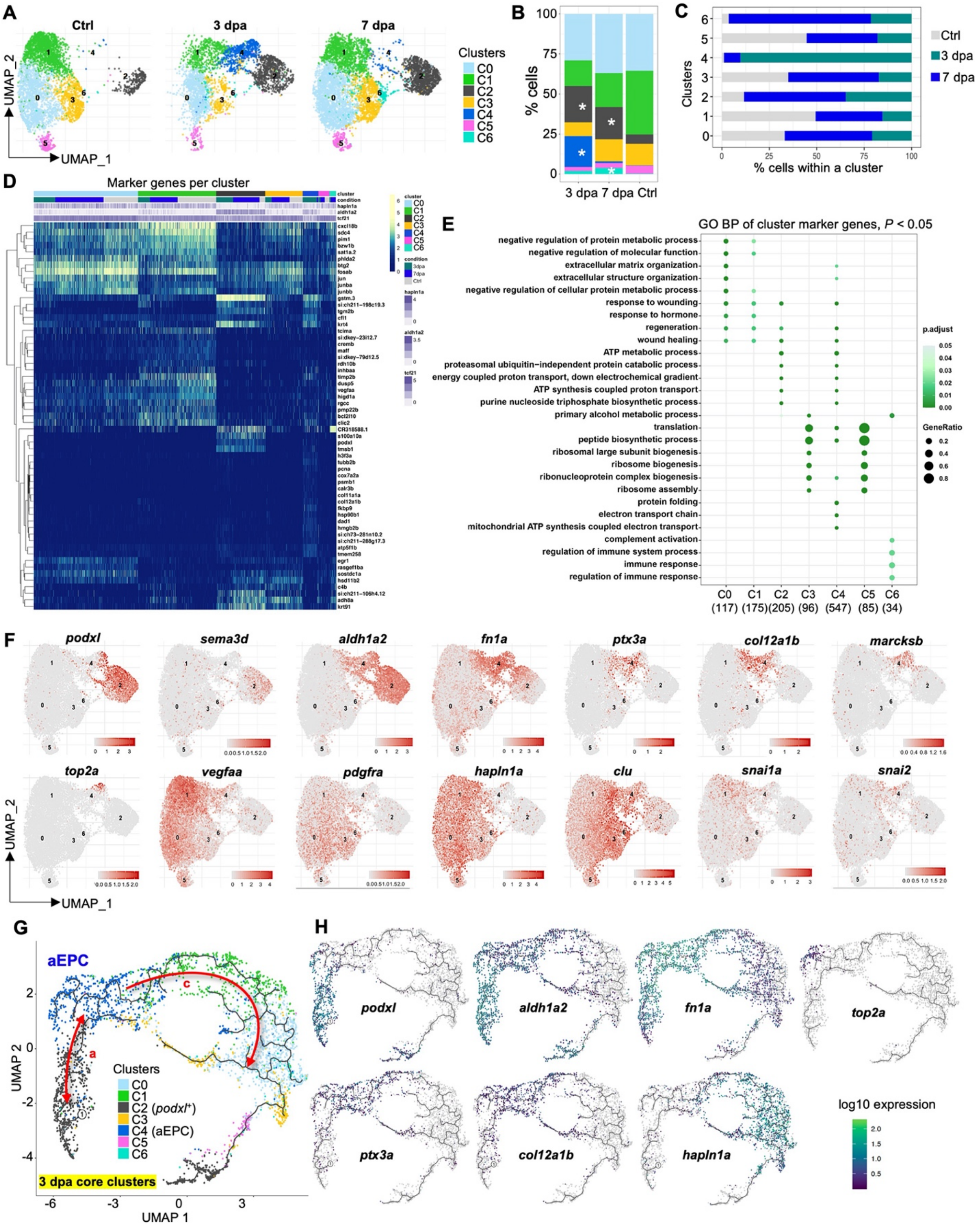


**Figure S3. Subsets of the core epicardial clusters.**

(**A**) UMAP of cell clusters of the core population across samples after re-clustering following the removal of mural cells and presumed contaminants. Cells are re-clustered as Core 1 (or C1) to C6, which are slightly different from the former clusters shown in Figure 2. The 3 and 7 dpa-enriched subsets are now clustered as C4 and C6, respectively. (**B, C**) Distribution of each sample's cells across the different core clusters. Cluster C2 expanded during regeneration, while clusters C0, C1, C3, C5, and C6 were reduced at 3 dpa before regaining relatively similar percentages as the uninjured control by 7 dpa. (**D**) Heatmap of top marker genes defining the clusters detected by both SC3 and Seurat. The order of the cells was determined by hierarchical clustering within each core cluster. *tcf21*, *aldh1a2*, and *hapln1a* expression are highlighted on top, but did not influence the cell ordering, which was driven by the genes within the heatmap. Normalized expression values are shown. (**E**) Biological process terms enriched for cluster marker genes in each cluster. The gene ratio is indicated by the dot size and the significance by the color of the dot ($P < 0.05$). The numbers of marker genes are listed in brackets under each cluster label. Notable GO terms include ECM organization (clusters C0 and C4), regeneration (C0, C1, C2, and C4), metabolic processes (C2 and C4), translation (C3, C4, and C5), and immune responses (C6). This suggests that the entire epicardial population actively participated in the regeneration process with diverse cellular and paracrine contributions. (**F**) Expression of selected marker genes on UMAPs (3 samples combined). (**G**) Pseudotime analysis results of the core epicardial clusters. Red arrows and letters highlight different trajectories. Clusters are labeled in the same number and color as in (A). (**H**) UMAPs showing expression patterns of genes correlated with the pseudotime trajectory.

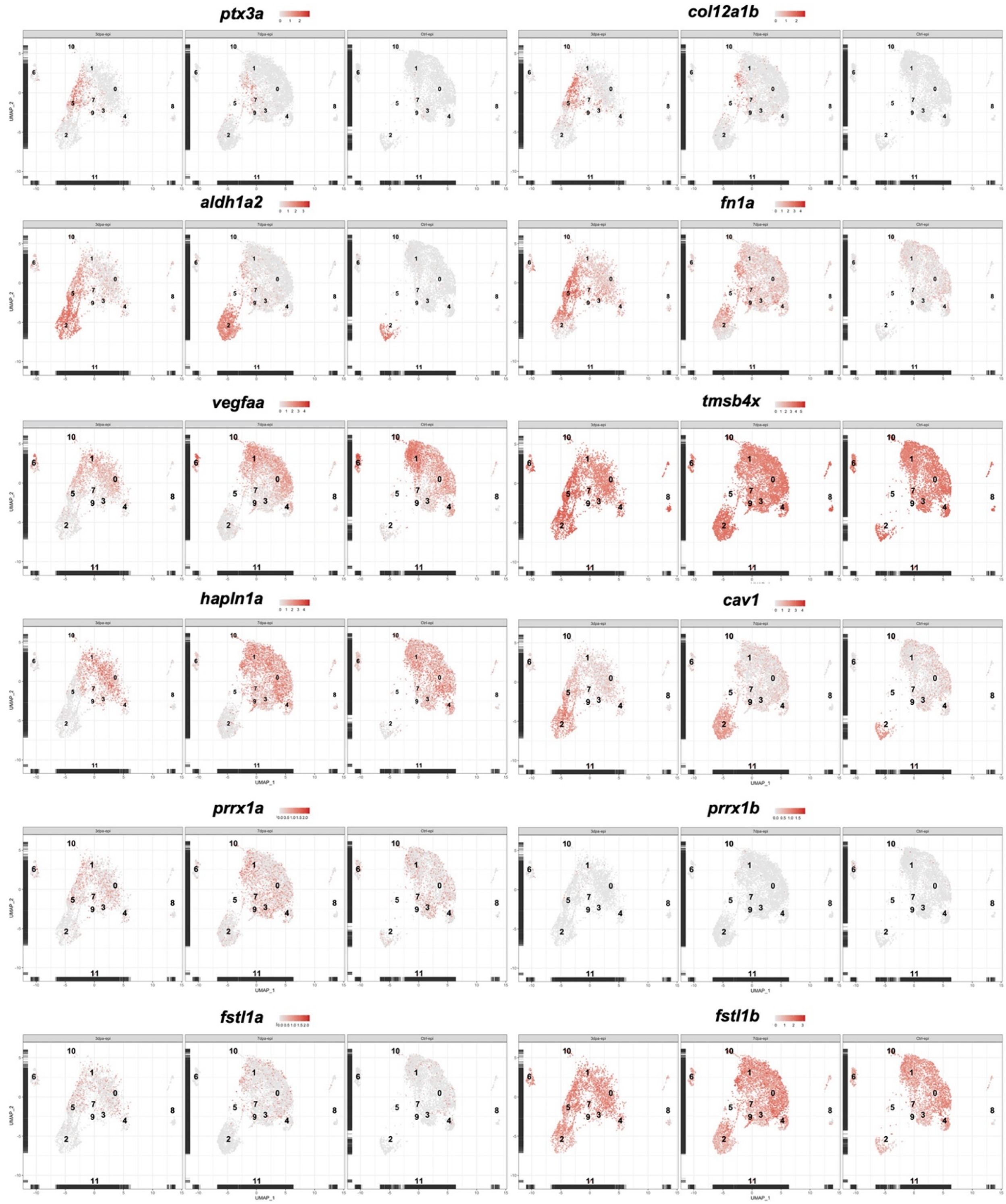


**Figure S4. Expression patterns of selected genes across all clusters and samples.**

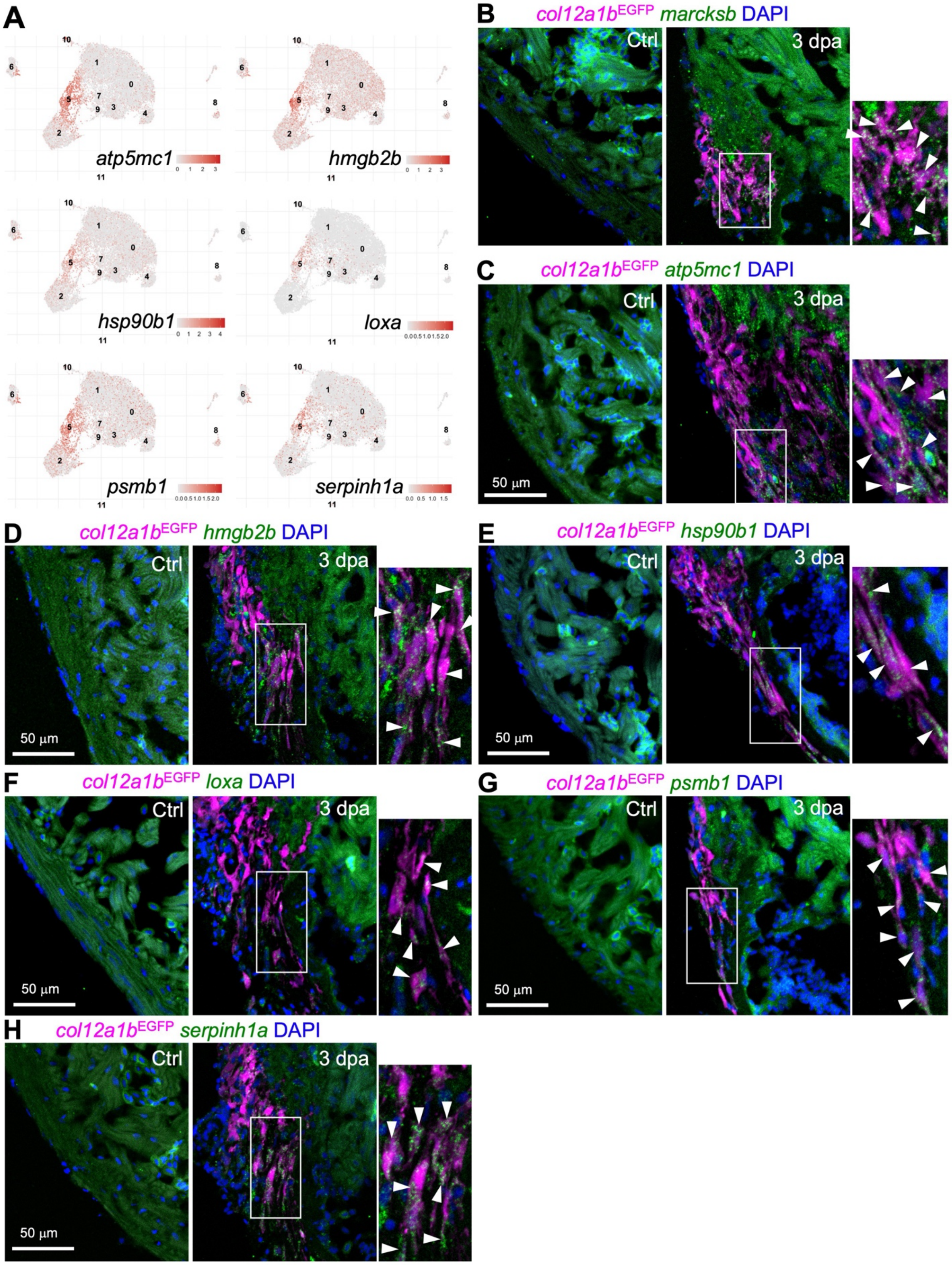


**Figure S5. Additional markers of aEPCs**

(**A**) UMAPs showing gene expressions. (**B-H**) Images of heart sections showing HCR staining results of selected aEPC markers in green, the *col12a1b*$^{EGFP}$ reporter in magenta, and DAPI staining in blue. The injury sites of 3 dpa samples and the equivalent regions of the uninjured hearts (Ctrl) are shown. The framed regions are enlarged to show on the right with arrowheads indicating representative HCR staining signals in EGFP$^{+}$ cells. Scale bar, 50 μm.

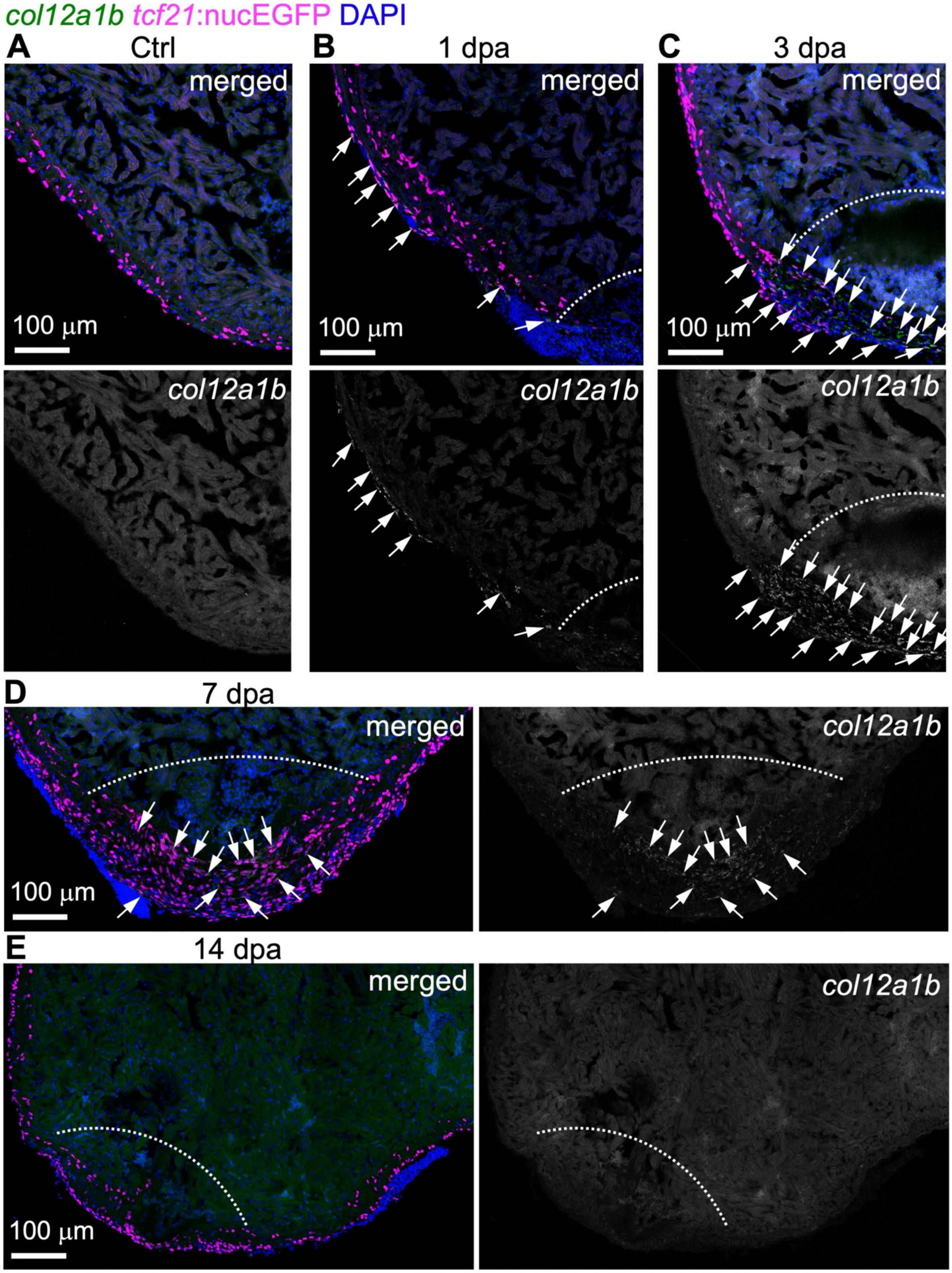


**Figure S6. *col12a1b* expression.**

HCR staining results of *col12a1b* (green) on heart sections collected at 1 (**B**), 3 (**C**), 7 (**D**), and 14 dpa (**E**) together with the uninjured control (**A**). *tcf21*:nucEGFP (magenta) labels the epicardial cells. Nuclei were stained with DAPI (blue). Single-channel images show signals of *col12a1b*. White dashed lines indicate the injury sites. Arrows denote representative *col12a1b*$^{+}$EGFP$^{+}$ cells. Scale bars, 100 μm.

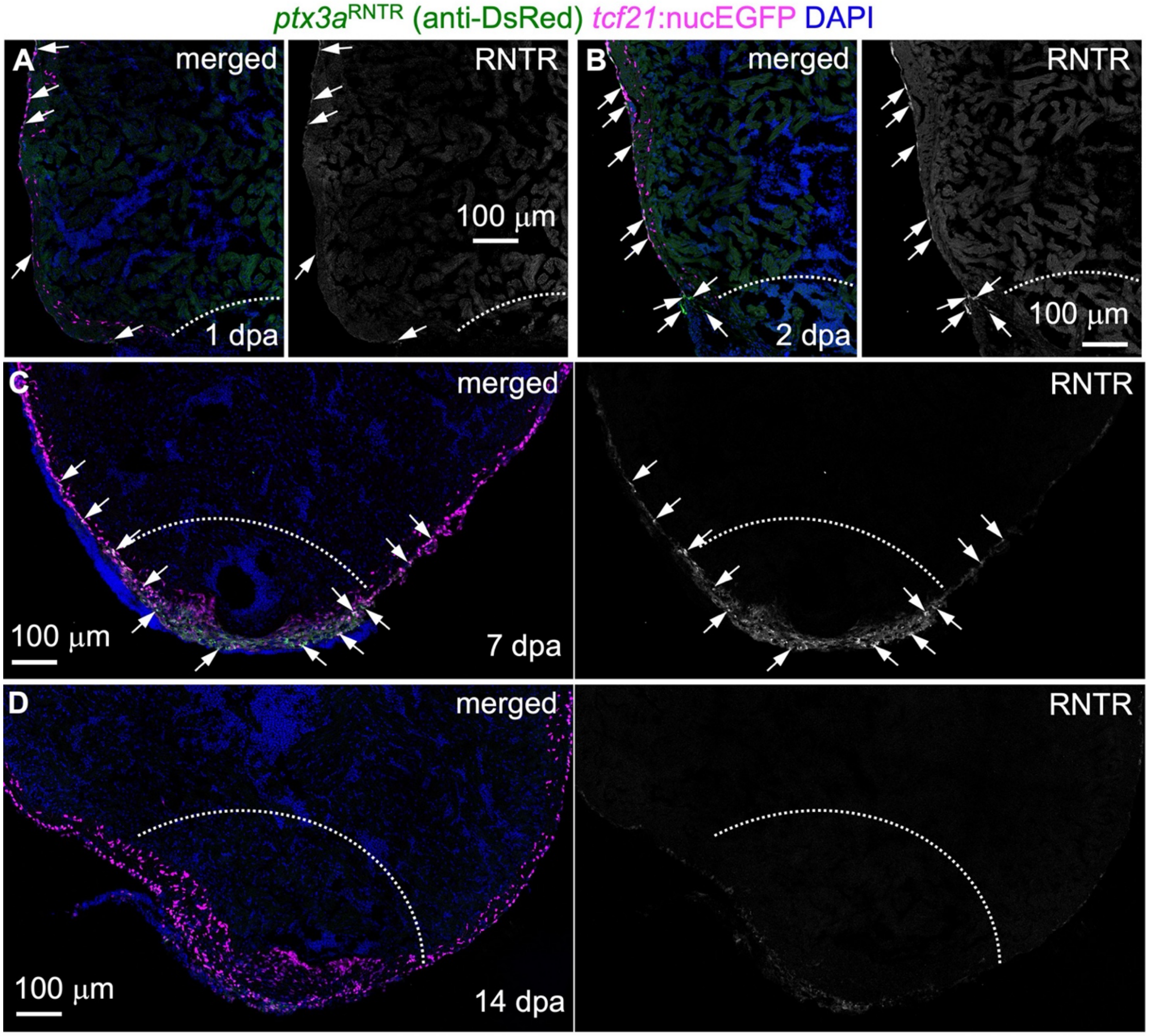


**Figure S7. Expression pattern of the *ptx3a*$^{RNTR}$ reporter line.**

Images of heart sections collected at 1 (**A**), 2 (**B**), 7 (**C**), and 14 dpa (**D**) showing expression of the *ptx3a*$^{RNTR}$ reporter in green. *tcf21*:nucEGFP (magenta) labels the epicardial cells. Nuclei were stained with DAPI (blue). Single-channel images show signals of *ptx3a*$^{\mathrm{RNTR}}$ (with anti-DsRed antibody staining) on the right. White dashed lines indicate the injury sites. Arrows denote representative RNTR$^{+}$EGFP$^{+}$ cells. Scale bars, 100 μm.

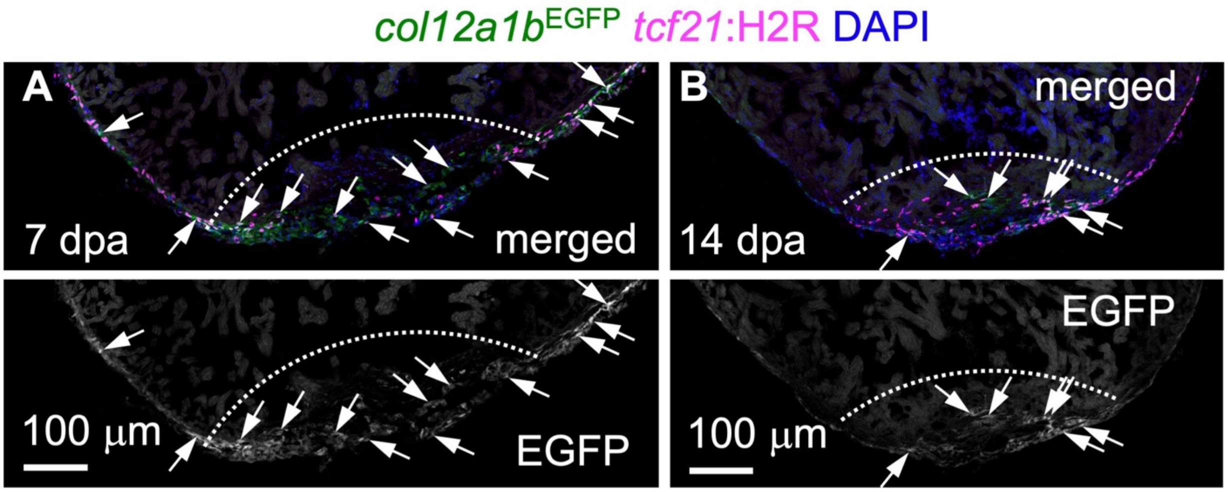


**Figure S8. Expression patterns of the *col12a1b$^{EGFP}$* reporter line.**

Images of heart sections collected at 7 (**A**) and 14 dpa (**B**) showing expression of the *col12a1b$^{EGFP}$* reporter in green. *tcf21*:H2R (magenta) labels the epicardial cells. Nuclei were stained with DAPI (blue). Single-channel images show signals of *col12a1b$^{EGFP}$*. White dashed lines indicate the injury sites. Arrows denote representative EGFP$^{+}$mCherry$^{+}$ cells. Scale bars, 100 μm.

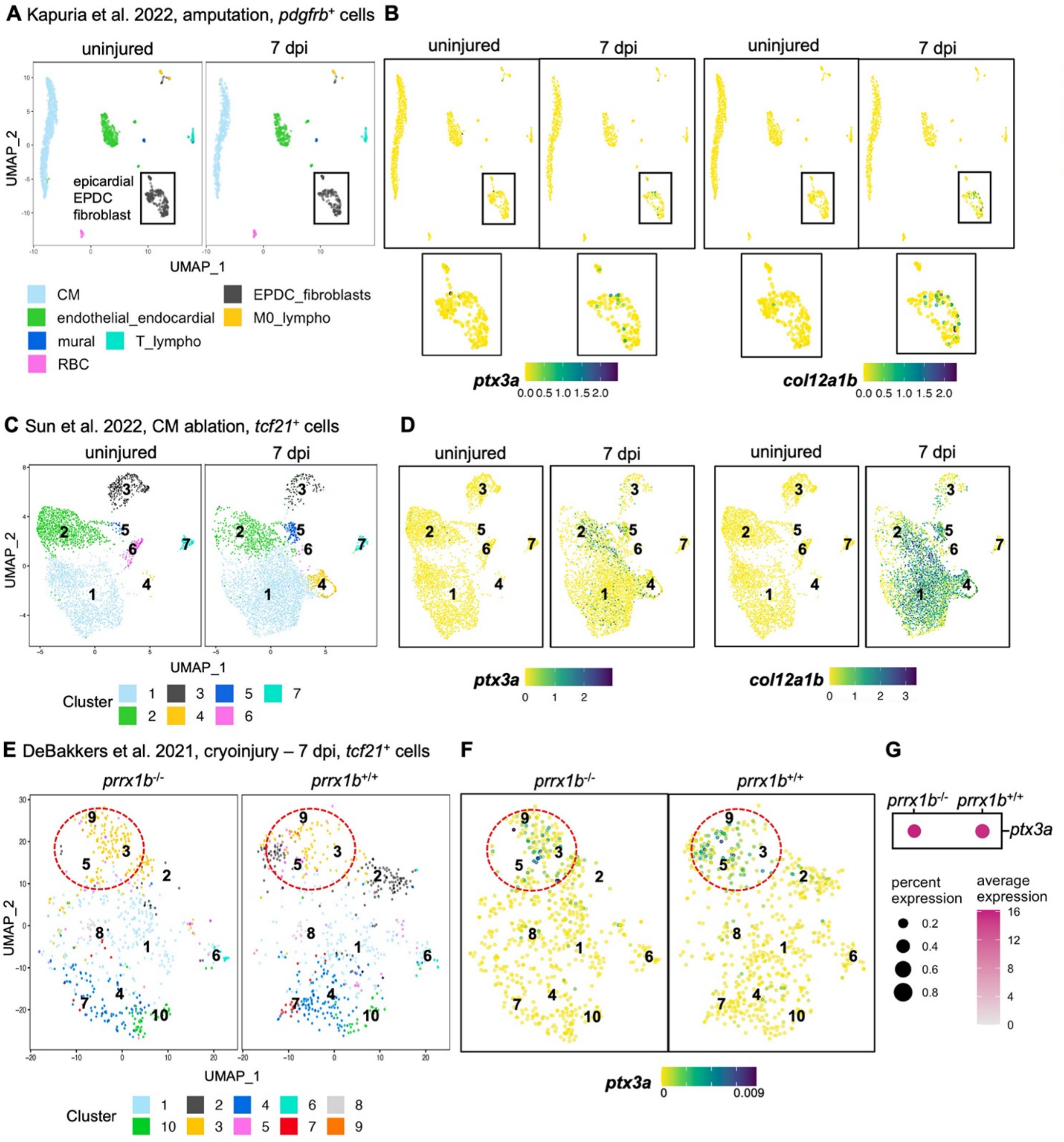


**Figure S9. Comparisons with other scRNA-seq datasets of zebrafish epicardium.** (**A-B**) UMAPs of the Kapuria et al. 2022 dataset (amputation injury, isolated *pdgfrb*$^+$ cells) showing cell clusters (A) and gene expressions (B)**.** Expressions of *ptx3a* and *col12a1b* are induced in the epicardial/epicardial-derived cell (EPDC)/fibroblast cluster at 7 days post-injury (dpi). (**C-D**) UMAPs of the Sun et al. 2022 dataset (CM ablation injury, isolated *tcf21*$^+$ cells) showing cell clusters (C) and gene expressions (D)**.** Expressions of *ptx3a*

and *col12a1b* are induced by injury 7 dpi. (**E-F**) UMAPs of the DeBakkers et al. 2021 dataset (cryoinjury, isolated *tcf21*$^{+}$ cells) showing cell clusters (E) and gene expressions (F) in the wild type (*prrx1b*$^{+/+}$) and *prrx1b*$^{-/-}$ samples. *ptx3a* is expressed in clusters 2, 3, 5, and 9. (**G**) Dot plot showing percentages of cells expressing *ptx3a* (41% in the mutant versus 47% in the wild type) and the average expression levels.

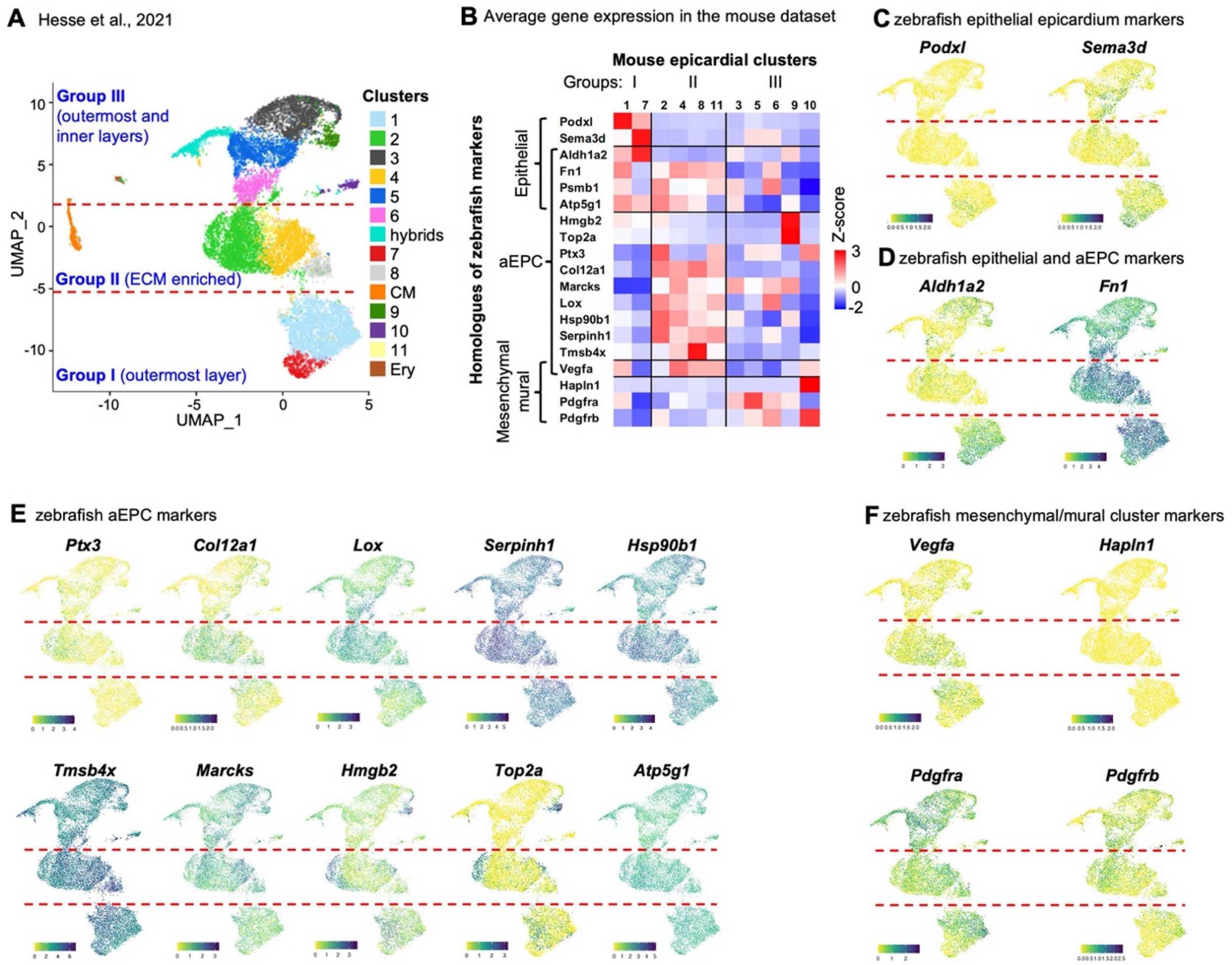


**Figure S10. Comparison with mouse epicardial stromal cells (EpiSC).**

(**A**) UMAPs of the Hesse et al. 2021 dataset showing cell clusters. Three groups of EpiSC were identified. Group I represents the outermost layer of the epicardium. Groups II and III cells reside in both the outermost and inner layers, and group II cells are enriched with expression ECM-related genes. (**B**) Heatmap showing average gene expressions of zebrafish epicardial cell markers in the mouse dataset. (**C-F**) UMAPs showing gene expressions of zebrafish epicardial cell markers in the mouse dataset.

**Supplementary Tables**

**Table S1.** List of marker genes for all clusters.

**Table S2.** List of marker genes for the core epicardial clusters.

**Table S3.** Enrichment annotation results of the marker genes for core epicardial clusters.
